# A geometric-to-neural cascade for cerebral microbleed detection in susceptibility-weighted MRI

**DOI:** 10.64898/2026.07.24.740624

**Authors:** Sam Bogdanov, Gaurav Rudravaram, Adam M. Saunders, Michael E. Kim, James LeFevre, Jack Charles, Stuti Jain, Matthew S. Schrag, Bennett A. Landman

## Abstract

Cerebral microbleeds (CMBs) are established imaging biomarkers of cerebral small vessel disease and are a defining feature of cerebral amyloid angiopathy (CAA), yet their automated detection in susceptibility-weighted imaging (SWI) remains challenging due to high false-positive rates from vessel cross-sections, iron and calcium deposits, and other hypointense mimics. We present a fully automated, three-stage cascade pipeline that combines subject-adaptive unsupervised candidate generation with two successive lightweight 3D ResNet classifiers, trained with only human-in-the-loop quality-assurance (QA) labels (yes/no per candidate) rather than dense voxel-wise segmentation masks. The candidate generation stage is performed by fitting a Gaussian Mixture Model (GMM) to each subject’s SWI intensity histogram to define an adaptive low-intensity threshold, followed by anatomical masking to exclude physiologically irrelevant regions (image edges, ventricles/CSF/choroid plexus, and cerebellum), and filters candidates by size and sphericity. The model was trained and evaluated across a nested 3×5-fold cross-validation on N = 30 subjects from a publicly available labeled microbleed dataset and a CAA cohort (11,424 CMB candidate lesions) with data augmentation during training. Stage A classifies all geometric candidates as CMB or non-CMB and Stage B refines the predicted positives to suppress false positives (cascade AUC = 0.9587, sensitivity = 0.712, specificity = 0.975, PPV = 0.676, F1 = 0.693). The cascade reduces Stage A false positives by 76.8% (888/1,157 false positives eliminated) while retaining competitive sensitivity. Inference was performed on 141 SWI scans, detecting a mean 40.3 CMBs per scan and being preferred for use in 85% of high CMB cases, as evaluated by a blinded neurologist. The inference pipeline outputs binary CMB segmentation NIfTI images and radiologist-ready QA visualizations.

## 1. INTRODUCTION

Cerebral microbleeds (CMBs) are small (2–10 mm in diameter) hypointense foci on susceptibility-weighted imaging (SWI) and T2*-weighted magnetic resonance imaging (MRI) arising from hemosiderin deposition following prior microhemorrhages [1]. They are established imaging biomarkers for cerebral small vessel disease, cerebral amyloid angiopathy (CAA), and hypertension-related arteriopathy [2], and are associated with elevated risk of stroke, cognitive decline, dementia, and mortality [3–6]. Manual CMB annotation is time-consuming, requires specialist neuroradiologists, and suffers from substantial inter-rater variability [7], motivating the development of reliable automated detection tools.

Several approaches to segmentation have been performed previously. Geometric-based automated methods relied on morphological features and intensity characteristics; one such example, the MAGIC model, was developed using Radon-based analysis, the outputs of which were fed into a random forest classifier. The MAGIC model was presented with a true positive rate (TPR, or sensitivity) of 0.95 but a positive predictive value (PPV, or precision) of 0.07-0.11 [7]. Parallel deep learning approaches have also been developed, primarily employing 3D convolutional neural networks (3D-CNN):

- Dou *et al*. [8] proposed a 3D-CNN for training that was then converted to a 3D fully connected network (FCN) for testing due to computational constraints (achieving TPR 0.93, PPV 0.44).
- Al-masni *et al*. [9, 10] combined SWI and Phase images as a two-channel input to facilitate a two-stage approach with YOLOv2 for candidate detection and a 3D-CNN for false-positive reduction (achieving TPR 0.71–0.86, PPV 0.62–0.67).
- Rashid *et al*. [11] introduced DEEPMIR, a 2D U-Net architecture trained on SWI and quantitative susceptibility map (QSM) image slices (achieving TPR 0.84–0.88, PPV 0.84).
- Sundaresan *et al*. [12] designed Microbleednet, which used geometric methods to generate candidate volumes, which were then supplied to a 3D U-net alongside the original image to generate whole-volume segmentations. The 3D U-net further served as a teacher model for a patch-based student model trained using knowledge distillation (achieving TPR 0.81–0.90, PPV 0.62-0.89).
- Lee *et al*. [13, 14] developed TPE-Det, an ensemble of axially-, sagittally-, and coronally-oriented 2D CNNs (achieving TPR 0.67, PPV 0.91).
- Tsuchida *et al*. [15] released SHIVA-CMB, a 3D U-net trained with Dice loss (achieving TPR 0.85–0.96, PPV 0.77–0.80).

Despite this progress, high false-positive (FP) rates in automated detection algorithms remain a core challenge because vessel cross-sections, iron deposits, and calcifications share the hypointense appearance of CMBs on SWI [16]. Threshold-based segmentation becomes difficult with fixed intensity thresholds due to scanner or acquisition protocol differences [1, 17]. Such issues are especially present when dealing with clinical data that are particularly heterogenous in their acquisition parameters. Moreover, obtaining ground-truth training labels requires extensive voxel-level manual annotation, limiting the availability of training data.

Publicly available, labeled datasets for training CMB detection algorithms must be carefully selected if they are to be used. While there exist several T2* Gradient Recalled Echo (GRE) datasets [15, 18–21], these scans suffer from poor through-plane resolution as well as reduced sensitivity when compared to SWI and are therefore clinically inferior to SWI for the detection of CMBs, particularly in CAA[22]. Furthermore, publicly available SWI datasets frequently present with a limited number of CMBs per scan (Dou *et al*. provide a dataset with 3.7 CMBs per scan [8]; Momeni *et al*. provide an SWI dataset with 2.25 CMBs per scan [18]). This makes the use of publicly available datasets challenging to train automated CMB detection algorithms for CAA. Consequently, publicly available datasets are of limited value, as T2*GRE scans are no longer the clinical standard [17, 22, 23] and existing SWI datasets with labeled images are not representative of CAA, which often presents with dozens to hundreds of CMBs per scan [22].

To circumvent these issues, we present a fully automated three-stage cascade pipeline trained on 11,424 patch labels using 10 SWI scans from an internal CAA clinical cohort and the dataset of 20 SWI scans provided by Dou *et al*. and outputs 3D segmentation masks. The pipeline comprises three components: (1) subject-adaptive unsupervised candidate generation combining a Gaussian Mixture Model (GMM) intensity threshold with geometric morphological filtering; (2) Stage A, a lightweight 3D ResNet trained to classify segmentation candidates as CMB or non-CMB with high sensitivity; and (3) Stage B, a second 3D ResNet that refines the predicted positives from Stage A, trained to discriminate true CMBs from FPs, increasing specificity. After training, we performed inference on 141 SWI scans in the CAA clinical cohort, where we found the proposed model detected more CMBs when compared to a recently developed publicly available model, SHIVA-CMB [15]. Our key contributions are subject-adaptive GMM thresholding generalizable across scanners, anatomical masking and geometric filters, and a two-phase cascade achieving specificity of 0.975 while retaining competitive sensitivity of 0.712 and PPV of 0.676 on the patch-level evaluation.

## 2. METHODS

### 2.1 Dataset and preprocessing

SWI data from N = 30 subjects were obtained from an internal CAA clinical cohort (N = 10, with paired T1 scans) and from the cohort presented by Dou *et al*. (N = 20) [8]. Ethical oversight of human imaging deidentification and use in the study was provided by the Vanderbilt University Medical Center Institutional Review Board and authorized under IRB# 180287. The preprocessing pipeline is comprised of five steps. First, each SWI volume was reoriented to right-anterior-superior conventions (RAS) using *nibabel* v5.3.2 [24] prior to resampling. Second, images were resampled to 1 mm^3^ isotropic voxel size. Third, ANTs [25] was used to rigidly register SWI to the T1 for alignment with existing SynthSeg [26] parcellations generated from the T1 images acquired in the same session. In the case that paired T1 images were not available, SynthSeg parcellations were performed on the SWI image. Fourth, brain mask extraction was performed on the registered SWI using SynthStrip [27]. Fifth, using *nibabel* v5.3.2, masked brain voxels were intensity-normalized to [0, 1] following percentile-based clipping to the 1st and 99th percentiles.

### 2.2 Brain parenchyma mask and candidate generation

The candidate generation pipeline proceeded in following steps:

#### 2.2.1: Anatomical exclusions

Brain tissue was defined as all voxels corresponding to the skull-stripped brain volume from preprocessing (brain_mask). Ventricles, cerebrospinal fluid (CSF), choroid plexus, and cerebellum (labels 4, 5, 7, 8, 14, 15, 24, 31, 43, 44, 46, 47, 63) were excluded from the candidate search region. Additionally, a 5-voxel erosion of the brain edge was applied following the MAGIC study methodology to exclude cortical surface artifacts [7]. Regions where candidates could be found – the parenchyma mask – were defined as the intersection of the eroded brain mask and the selected SynthSeg parcellations. Figure 1 visualizes this masking process.

**Figure 1.**
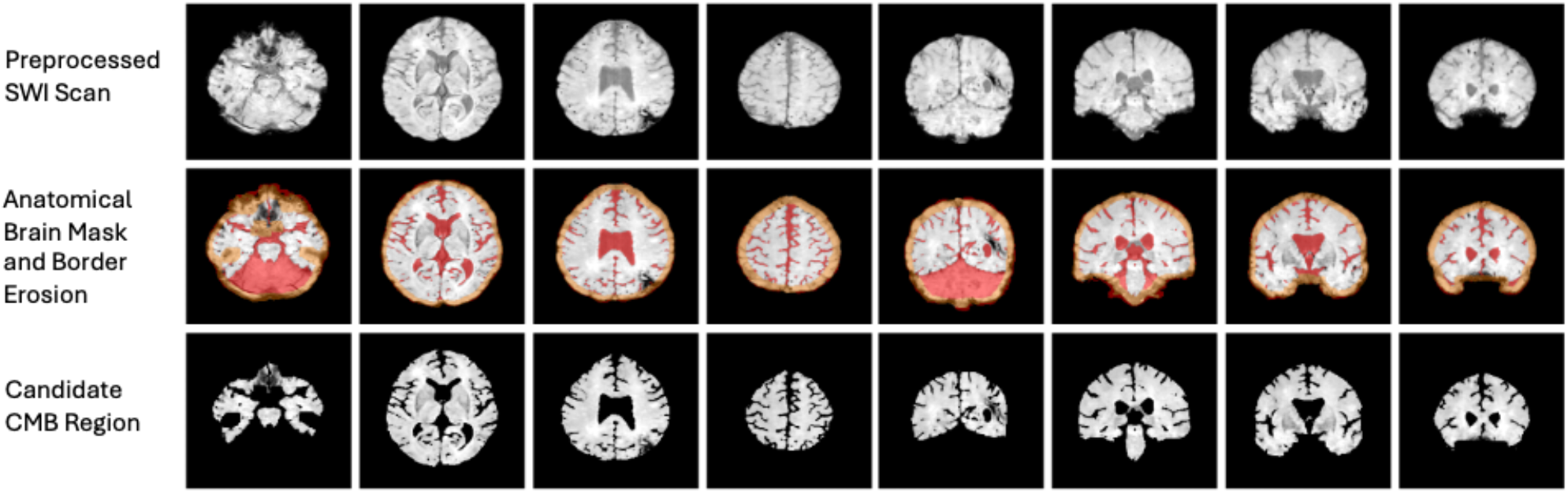
Brain parenchyma mask construction for one representative subject. Top: preprocessed SWI. Middle: excluded regions (red = SynthSeg-excluded ventricles/CSF/choroid plexus/cerebellum; orange = eroded 5-voxel brain border). Bottom: final parenchyma mask used for candidate search. Excluding these regions removes the tissue types most prone to susceptibility artifacts that mimic CMBs, so downstream candidate generation only searches genuine brain parenchyma.

#### 2.2.2: Gaussian Mixture Model (GMM) tissue segmentation

A 3-component GMM was fit to the normalized intensity histogram of each subject’s skull-stripped SWI. Initialization used the 10^th^, 35^th^ and 75^th^ percentiles as starting means to promote stable convergence following typical grey- and white-matter intensity distributions (Figure 2). Components were assigned as follows: the component with the largest standard deviation was assigned as background, then the lowest remaining mean as gray matter (GM) and the highest as white matter (WM). Per-subject parameters (μ, σ, weight) for GM were used to define a subject-specific low-intensity threshold.

**Figure 2.**
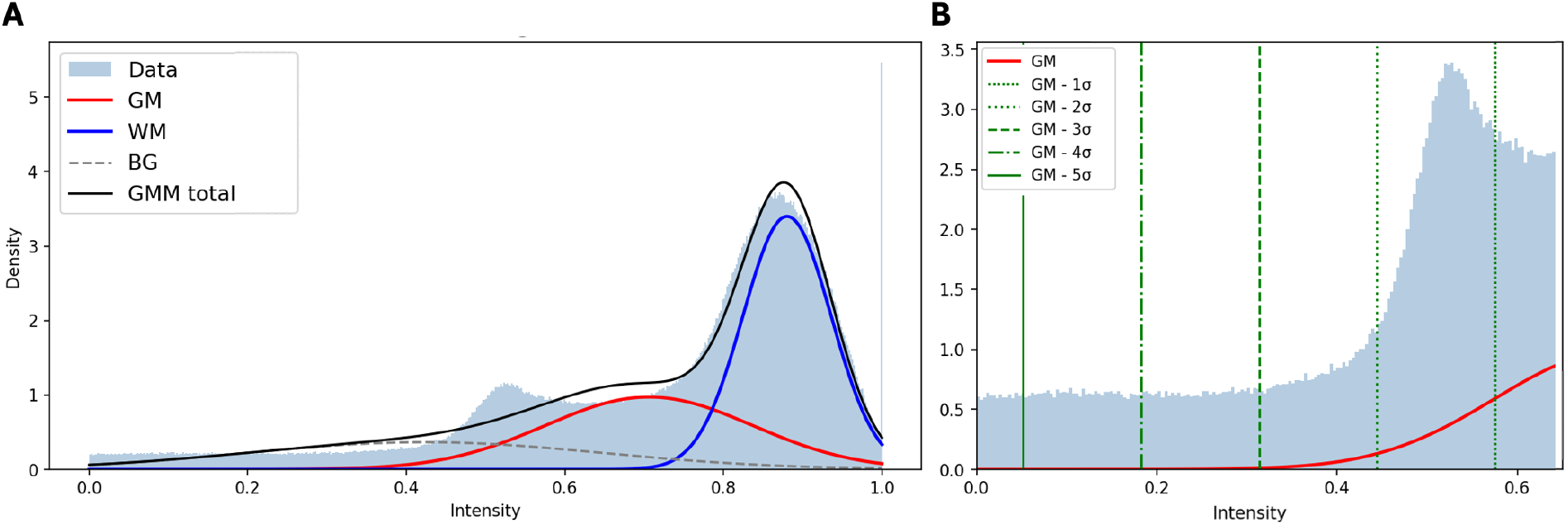
A 3-component Gaussian Mixture Model (GMM) fit to the SWI intensity histogram for a representative subject. GM = grey matter; WM = white matter; BG = background. A: full intensity range showing the three fitted components. B: low-intensity zoom with candidate thresholds at μ_GM_-1σ through μ_GM_-5σ overlaid; μ_GM_-1σ (the rightmost dotted line) was the operating threshold used for candidate generation, selected empirically as the point that captured the greatest number of true microbleeds while keeping the candidate search space, and therefore false-positive burden, manageable for downstream QA and classification.

#### 2.2.3: Geometric filtering

Hypointense voxels were identified per 2.2.2. Candidate components were then generated using diagonal-constrained priority region growing. Specifically, seeds were processed from darkest to brightest normalized SWI intensity, and each seed was expanded greedily through 26-connected hypointense neighboring voxels. During growth, a candidate was allowed to expand only if the Euclidean diagonal of its bounding box remained less than 10 mm, using the distance between the extreme voxel coordinates along each axis. After each component was finalized, a 1-voxel 26-neighbor buffer was marked around it to prevent immediately adjacent voxels from seeding a separate candidate. Raw candidates were then filtered geometrically: components with volume less than 2 voxels, bounding-box diagonal greater than 10 mm, or aspect ratio greater than 3.0 were rejected. The aspect-ratio filter removed elongated structures, such as vessels or sulcal blood products, by rejecting candidates whose longest bounding-box axis was more than three times the shortest axis.

After visual inspection, 1 standard deviation below the GM mean was chosen as the standard cutoff since it captured the most CMBs while still producing fewer than 2,000 candidates per scan for manual QA.

### 2.3 Human-in-the-loop label generation

Manual quality assurance was performed on geometric candidates from the 10 CAA subjects using a custom review interface by Kim *et al*. [28] displaying three views per candidate: (1) the candidate mask overlaid on SWI (zoomed), (2) the raw SWI patch at the candidate location, and (3) the full axial slice with bounding box annotation. Candidates were labeled “Yes” (confirmed CMB) or “No” (FP mimic). For the 20 scans with ground-truth labels from the dataset generated by Dou *et al*., a “Yes” label was assigned if the candidate CMB contained the ground-truth positive coordinate and “No” if the candidate did not.

The final dataset comprised 11,424 candidates for QA across 30 scans (8,313 from 10 CAA scans, 3,111 from 20 scans from the dataset supplied by Dou *et al*.). All scans came from independent patients, with 787 positives (6.9%) and 10,637 negatives. Per-subject label counts are reported in Supplemental Table S2. Candidate patches per participant were not equally distributed, instead relying strictly only on candidates detected via GMM and geometric correction.

For each QA-labeled candidate, a 16×16×16 voxel patch was extracted from the preprocessed SWI volume centered on the candidate centroid in voxel coordinates. Patches exceeding image boundaries were zero-padded. Each patch was stored alongside metadata including subject identifier, session, SWI path, centroid coordinates, and binary label.

### 2.4 Stage A: CMB vs. Background 3D ResNet

#### 2.4.1: Architecture

The ResNet3DClassifier is a compact 3D residual network inspired by He *et al*. [29] with modifications to ResNet depth and model size for 16 × 16 × 16 volumetric patches. The model contains 661,041 trainable parameters and uses three residual stages with two residual blocks per stage. Channel depth increases from 16 → 32 → 64 across stages, while spatial resolution is halved after each stage from 16^3^ → 8^3^ → 4^3^ → 2^3^. The resulting 64-channel 2 × 2 × 2 feature map is collapsed by global average pooling into a 64-dimensional vector, followed by a final linear layer that produces a single output logit. The network design can be seen in the Figure 3 subpanel.

**Figure 3.**
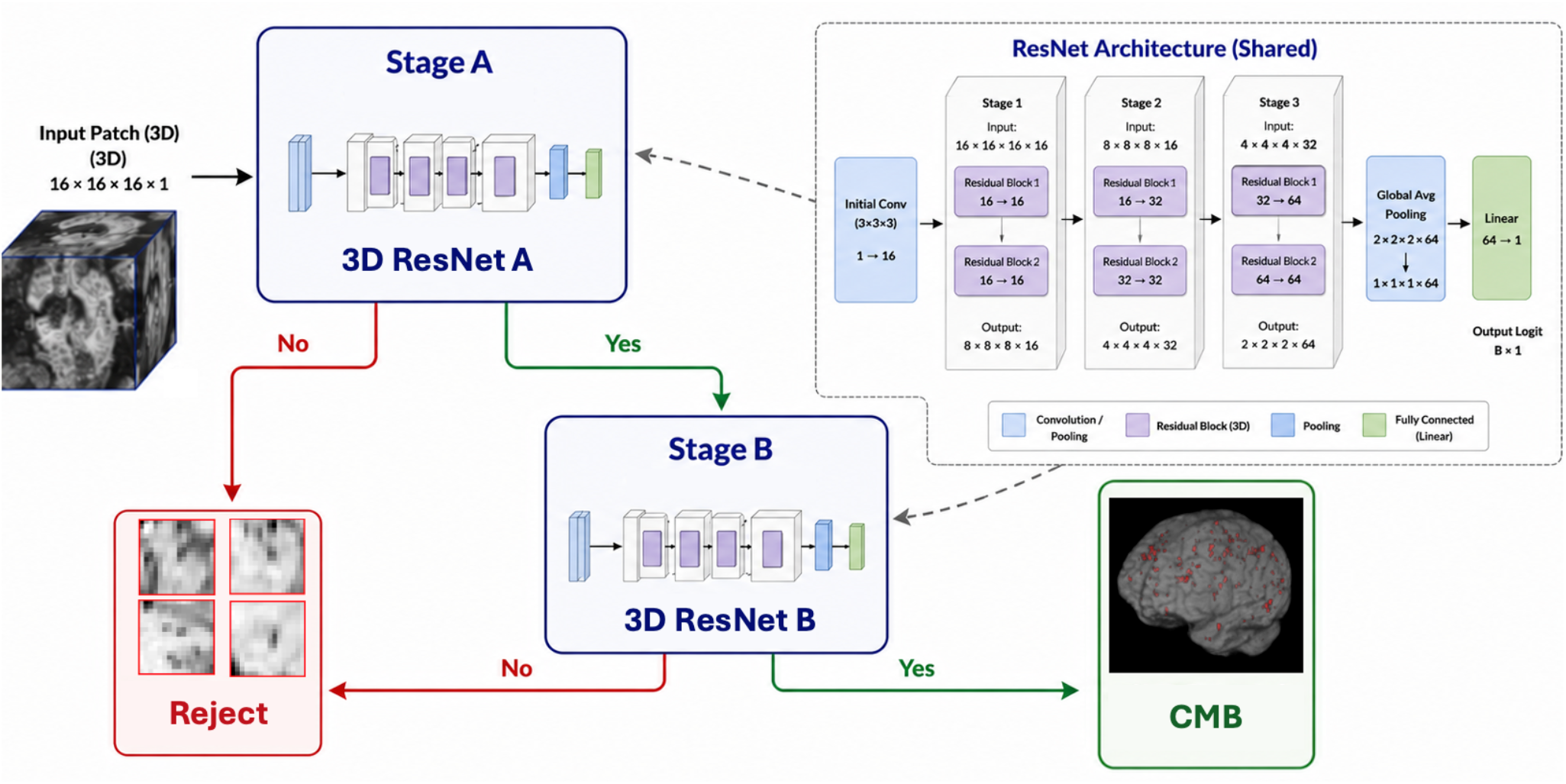
Cascade flow through 3D ResNets. The architecture presented in the subpanel was used identically for both Stage A and Stage B classification. The lightweight residual design (661K, parameters, adapted from He *et al*. 2016) processes 16^3^-voxel patches through three stages of channel-doubling and spatial down sampling. ResNets are sized for efficient inference across thousands of candidates per cohort. Predicted positives from Stage A are fed into Stage B, where they are confirmed to increase model specificity.

#### 2.4.2: Training protocol

A nested 3-outer x 5-inner fold cross-validation was performed, with splits assigned at the subject level and proportional representation from the CAA and Dou datasets in each split. The loss function was binary cross entropy (BCE) with positive weight (n_neg_/n_pos_) specific to each training fold (range 7.12 – 32.44, average 13.51 across all GT labels) to address class imbalance in gradient magnitude, supplemented by weighted random sampling by n_neg_/n_pos_ to achieve expected equal positive and negative patch presentation during training. This redundancy was chosen to promote accurate detection of positive classes. Optimization used Adam [30] with learning rate 10^-3^, and a reduced learning rate scheduler (mode = min, patience = 10, factor = 0.5). Models were trained for 100 epochs per fold, with the checkpoint at minimum validation loss retained. Data augmentation applied during training included random axis-wise flips (p = 0.5 per axis), random 90° axial-plane rotation (uniform from {0°, 90°, 180°, 270°}), random intensity scaling (from [0.9, 1.1]), and random intensity shift (from[-0.05, 0.05]).

### 2.5 Stage B: Refinement of Predicted Positives

#### 2.5.1: Dataset construction

For each outer fold, Stage A predictions from all five inner-fold models were aggregated by computing the mean probability per patch. Patches with mean probability greater than or equal to 0.5 were forwarded to Stage B. This yielded 1,890 patches (733 true CMBs, 1,157 FPs).

#### 2.5.2: Training protocol

Stage B uses an identical ResNet3D classifier to Stage A. Training matched what was performed in Stage A. A hyperparameter comparison over learning rate ∈ {10^-3^, 10^-4^} and dropout ∈ {0, 0.3} was conducted on cascade performance; learning rate = 10^-3^ with no dropout was selected given its best performance across all metrics (Table 1).

**Table 1.** Stage B cascade performance across a hyperparameter grid search (learning rate × dropout) on CAA-only data. The best configuration (learning rate = 10^−3^, no dropout) was retrained on the combined CAA + Dou dataset (bottom row) to produce the final reported cascade model.

| Dataset | Learning Rate | Dropout | Cascade AUC | Sensitivity (TPR) | Specificity | PPV (Precision) | F1 |
| --- | --- | --- | --- | --- | --- | --- | --- |
| CAA | $10^{-3}$ | 0 | 0.9502 | 0.671 | 0.963 | 0.731 | 0.700 |
| CAA | $10^{-3}$ | 0.3 | 0.9490 | 0.662 | 0.963 | 0.729 | 0.694 |
| CAA | $10^{-4}$ | 0 | 0.9430 | 0.602 | 0.960 | 0.697 | 0.646 |
| CAA | $10^{-4}$ | 0.3 | 0.9422 | 0.595 | 0.959 | 0.687 | 0.638 |
| CAA + Dou | $10^{-3}$ | 0 | 0.9587 | 0.712 | 0.975 | 0.676 | 0.693 |

#### 2.5.3: Cascade inference

A patch is classified as a CMB if (1) Stage A probability ≥0.5 and (2) Stage B probability ≥0.5. The two-stage design explicitly trades sensitivity for specificity: Stage A casts a wide net while Stage B prunes FPs. The network design and cascade flow can be seen in Figure 3.

### 2.6 Cascade Network Performance

Table 1 reports cascade performance across all four hyperparameter configurations, evaluated on the CAA dataset only. The selected configuration (learning rate = 10^-3^, no dropout) yields cascade area under the receiver-operator curve (AUC) = 0.959, sensitivity = 0.712, specificity = 0.975, PPV = 0.676, and F1 = 0.693, corresponding to TP = 560, FN = 227, FP = 269, TN = 10,368. Compared to Stage A alone (specificity = 0.891, sensitivity = 0.931), the cascade reduces the FP count from 1,157 to 269, representing a 76.8% reduction, at the cost of 173 additional FNs (FN increase from 54 to 227). TPs are reduced by 23.6%, from 733 to 560. Figure 4 illustrates the side-by-side confusion matrices.

**Figure 4.**
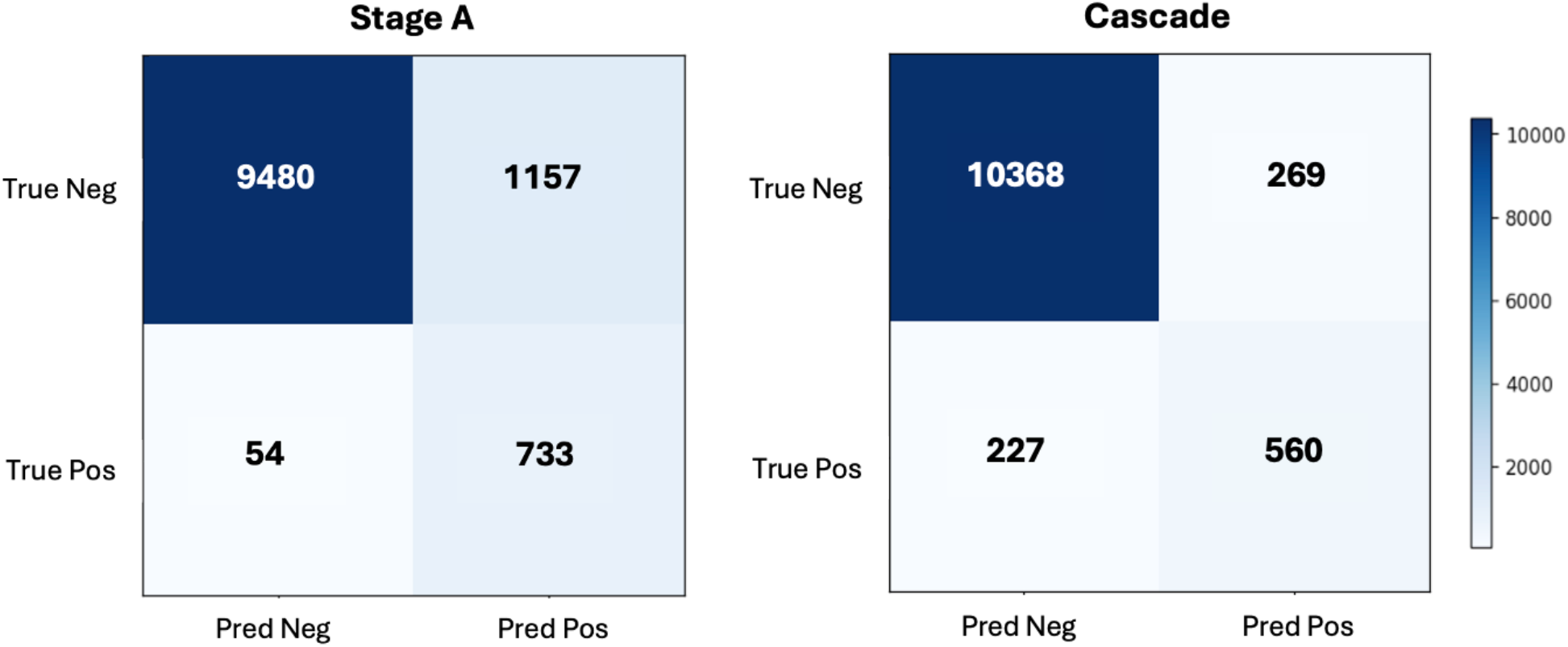
Comparison of Stage A standalone vs. full cascade (A → B). The cascade substantially reduces FPs (1157 to 269) while accepting reduced sensitivity (0.931 to 0.712).

**Figure 5.**
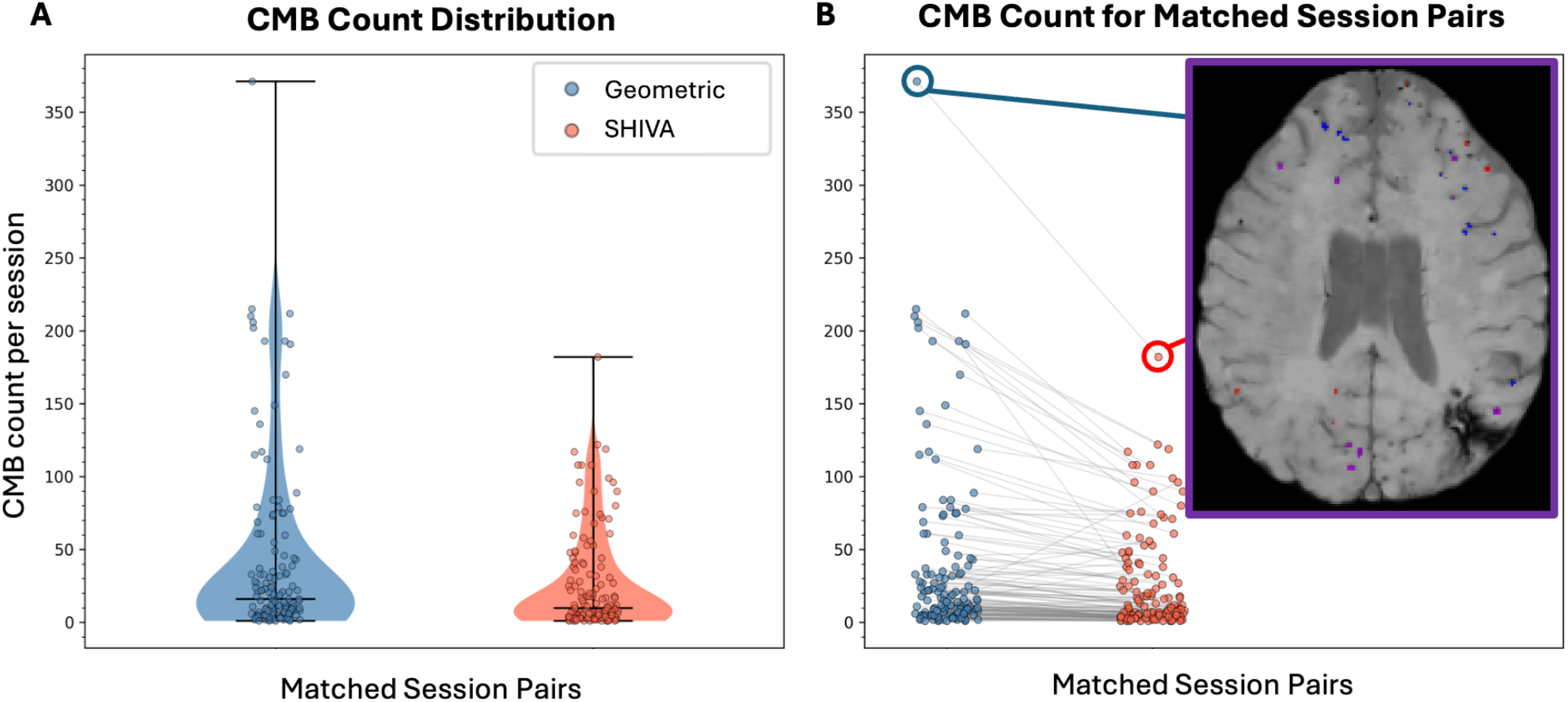
CMB detection algorithm (Geometric) compared to SHIVA-CMB in a CAA cohort. A: CMB count distribution per algorithm. B: paired counts per matched session; inset shows segmentation overlaps for the highest-burden session (purple = agreement, blue/red = Geometric-/SHIVA-only). Geometric detects substantially more CMBs than SHIVA in high-burden sessions, and comparison of the non-overlapping (blue) detections to the underlying scan supports that these are true CMBs.

### 2.7 Inference pipeline

The deployed inference pipeline executes three phases for each input subject and session. Phase 1 performs geometric candidate generation: preprocessing, GMM fitting, threshold application, sphericity and size filtering, and SynthSeg exclusion (preferentially employing a candidate NIfTI with paired T1 image, but applicable to SWI images alone). Phase 2 runs Stage A classification: patch extraction followed by ResNet ensemble prediction (aggregated across trained inner-fold models), retaining patches with mean probability ≥0.5. Phase 3 runs Stage B classification on the retained patches, keeping those with mean probability ≥0.5 as final CMB detections in a segmentation file.

## 3. RESULTS

### 3.1 Comparison to prior work

We sought to compare the proposed algorithm to the highest-performing publicly available model, SHIVA-CMB [15]. Due to a lack of publicly available datasets on which SHIVA-CMB was not trained, we chose to evaluate the proposed algorithm against SHIVA-CMB on 20 of the highest burden scans from the CAA clinical cohort, as defined by number of CMBs detected via the proposed method. The segmentations were presented side-by-side, along with the baseline, unlabeled image. A trained neurologist with over 10 years of experience and several hundred CAA patients treated (author MSS) was blinded to the algorithms used to generate each segmentation and rated the segmentations on their performance, preferring the use of the proposed CMB detection algorithm to SHIVA in 17 of 20 cases.

### 3.2 Inference on CAA cohort

The CMB detection pipeline was deployed on 141 SWI scans in the CAA clinical cohort (excluding minimal intensity projection images because of anisotropy). The detection algorithm was robust across 29 SWI acquisition protocols. 138 scans (97.9%) had 1 or more detected CMB, with a mean of 40.3 CMB, a standard deviation of 59.6, and median of 16 CMB for all scans. When compared to SHIVA-CMB inference scans run on the same scans, the proposed algorithm detected significantly more CMBs per scan for pairs of scans with ≥1 CMB detected per scan (p < 0.05, pairwise Wilcoxon signed-rank test, n = 132).

## 4. DISCUSSION

The proposed cascade approach was robust to the detection of CMB in the CAA cohort, a notably challenging cohort on which previous segmentation approaches (MAGIC, SHIVA-CMB, and Microbleednet) had underperformed. We attribute the performance of the proposed model to the minimal introduction of scan-wide contrast differences by focusing on patches instead of whole image volumes, which can improve generalizability; similarly, previous non-machine-learning approaches were devised on single-site images [7], limiting their robustness. The patch data were sourced from 11 different sites/acquisition parameters (10 from an internal CAA clinical cohort, 1 from Dou *et al*.) and data were further augmented with intensity scaling and intensity shift, enabling for robust out-of-domain generalization.

The cascade design separates two complementary objectives: high recall in Stage A and high precision in Stage B. The explicit probability threshold at each stage allows the sensitivity/specificity tradeoff to be tuned for clinical use without retraining. Subject-adaptive GMM thresholding avoids hardcoded intensity cutoffs that fail across scanner types and acquisition protocols. SynthSeg parcellations were resampled into SWI space using each image’s affine transform rather than its array dimensions, since subjects’ parcellation and SWI volumes could share the same voxel grid size but have different affines.

Stage A achieved strong recall (sensitivity 0.931) with a patch-level AUC of 0.9673, serving as an effective high-recall screen. Stage B successfully filtered 76.8% of Stage A false positives with a relatively modest reduction (23.6%) in true positives. The learning rate = 10^-3^ configuration without dropout consistently outperformed alternatives at the cascade level, consistent with the near-balanced Stage B training set reducing the need for regularization. A patch-level F1 of 0.693 is comparable to prior work on CMB segmentation.

The model presented in this work detects more microbleeds compared to SHIVA-CMB [15] – the publicly-available model with the highest published performance metrics – in a cohort of clinical scans from patients with CAA (p = 6.095 × 10^-14^), demonstrating its utility in imaging for this specific disease. Visualization of CMB counts from SHIVA in Figure 4 suggest that the model has a strong preference to keep CMB counts less than 100, and absolutely under 200; we hypothesize that this may be the result of using whole image segmentation and training on a dataset which was not as burdensome as the internal CAA clinical cohort data, thereby limiting total detection capacity. In turn, the proposed model considers each patch independently, so a global limit is not learned. Direct comparison of the proposed approach to SHIVA-CMB is challenging because we could not find any publicly available, ground truth labeled SWI datasets which had not been used in the process of training SHIVA-CMB and we did not have densely labeled segmentation masks. Identification of such a suitable test set would allow for quantitative, direct comparison of the models.

The training cohort is moderate (N=30 subjects) but diversified in its acquisition types (N=11 acquisition parameters). The addition of more scans and acquisition types is likely to improve model performance, but it is unclear if there is a limit on the reduction in sensitivity for the cascade model given the model architecture; we hypothesize that an increase in the amount of training data for the Stage B model could overcome the drawback of decreased sensitivity when implementing the cascade model. However, this potential bound on sensitivity is offset by the increased detection capacity for high-burden scans and the model’s robustness in the face of variable acquisition types, as demonstrated when deployed across 141 heterogeneously acquired SWI scans (29 acquisition types) from the CAA cohort.

The use of a cascade model to deliver moderate sensitivity (TPR) with high specificity of this tool is of utility for clinicians and researchers, providing high-throughput quantification; although CMBs are being missed by the proposed method, they can readily be manually augmented while the high specificity of the model ensures strong applicability in the clinical setting, and we further provide tools to perform quality-control for FPs. The development of such a manual augmentation tool for the refinement of detections in the CAA cohort can enable improved training of an identical two-phase model with a complete dataset or full-volume segmentation using nn-UNet or novel 3D U-Net structures. Accordingly, the results from the proposed method pave the way for future work investigating complementary strategies, including scribble-supervised (partial cross-entropy) training objectives to better utilize spaces not detected as candidate patches, self-supervised pretraining on unlabeled SWI data, and uncertainty-aware ensemble aggregation, to further improve model sensitivity while preserving the generalizability demonstrated by the proposed approach.

## 5. CONCLUSION

We present a three-stage cascade for automated CMB detection in SWI that combines subject-adaptive unsupervised candidate generation with two successive 3D ResNet classifiers. The pipeline was trained on only weak yes/no QA labels, achieved cascade AUC = 0.9587, sensitivity of 0.712, specificity of 0.975, and was fully deployed for inference on a CAA neuroimaging cohort. On this cohort, it outperforms existing methods in its detection of high lesion burdens. Our work provides a foundation for future integration of scribble-supervised losses and full-volume nnU-Net segmentation to further improve sensitivity of CMB detection.

## ACKNOWLEDGEMENTS

Research reported in this publication was supported by NIGMS of the National Institutes of Health under award number T32GM007347 and T32GM152284.

This work was supported by the Alzheimer’s Disease Sequencing Project Phenotype Harmonization Consortium (ADSP-PHC) that is funded by NIA (U24 AG074855, U01 AG068057 and R01 AG059716).

This work was supported by NIA of the National Institutes of Health under award number T32-AG058524.

Author Jack Charles is the 2026-2027 recipient of the Mary Frances Gray Lundy Endowed Scholarship for the DeBusk College of Osteopathic Medicine in memory of James Charles Gray, Sr.

During the preparation of this work the authors used ChatGPT, an AI language model developed by OpenAI, Gemini, an AI language model developed by Google, and Claude, an AI language model developed by Anthropic in order to assist in coding, image generation, text production, and text editing. After using these tools, the authors reviewed and edited the content as needed and take full responsibility for the content of the publication.

